# King- and queen-specific degradation of uric acid is the key to reproduction in termites

**DOI:** 10.1101/2022.09.28.510009

**Authors:** Takao Konishi, Eisuke Tasaki, Mamoru Takata, Kenji Matsuura

## Abstract

Caste-based reproductive division of labour in social insects is built on asymmetries in resource allocation within colonies. Kings and queens dominantly consume limited resources for reproduction, while non-reproductive castes such as workers and soldiers help reproductive castes. Studying the regulation of such asymmetries in resource allocation is crucial for understanding the maintenance of sociality in insects, although the molecular background is poorly understood. We focused on uric acid, which is reserved and utilized as a valuable nitrogen source in wood-eating termites. We found that king- and queen-specific degradation of uric acid is the key to reproduction in the subterranean termite *Reticulitermes speratus*. The urate oxidase gene (*RsUAOX*), which catalyses the first step of nitrogen recycling from stored uric acid, was highly expressed in mature kings and queens, and upregulated with differentiation into neotenic kings/queens. Suppression of uric acid degradation decreased the number of eggs laid per queen. Uric acid was shown to be provided by workers to reproductive castes. Our results suggest that the capacity to use nitrogen, which is essential for the protein synthesis required for reproduction, contributes to colony cohesion expressed as the reproductive monopoly held by kings and queens.

## 1. Introduction

Caste-based reproductive division of labour is a prominent characteristic of social insects; only a small fraction of individuals monopolize reproduction, where the others conduct non-reproductive tasks [1,2]. Such social systems are built on asymmetries of resource allocation within colonies. In many social insects, such as ants, bees, wasps and termites, rates of nutrient flow to larval and adult reproductive castes (kings and queens) are higher than to non-reproductive castes, such as workers and soldiers [3,4]. For example, in honeybees, reproductive/non-reproductive caste designation is determined through early nutritional exposure to “royal jelly” [5]. Larvae provided with a diet composed primarily of royal jelly grow more rapidly, have well-developed ovaries and emerge as queen bees. In contrast, worker bees are not provided with high levels of this nutrient-rich resource, and are therefore smaller with only rudimentary ovaries. Studying the regulation of such asymmetries in resource allocation is crucial for understanding the maintenance of sociality in insects; at present, the molecular background is not well-known. The capacity for nitrogen use, which is essential for the protein synthesis required for reproduction, can especially be hypothesized to be involved in colony cohesion expressed as the reproductive monopoly held by kings and queens.

Wood-eating termites feed on a nitrogen-poor diet and have evolved efficient nitrogen conservation strategies [6,7]. Uric acid is reserved and utilized as a valuable nitrogen source for termites [8,9]. As in a number of cockroach species, uric acid is a major product of purine metabolism stored in specialized urocyto cells of fat bodies in termites [10,11]. Termites are capable of transporting uric acid stored in fat bodies to the hindgut via the Malpighian tubules, whereas intestinal anaerobic bacteria efficiently recycle nitrogen via ammonia production and downstream amino acid synthesis [8,12–15]. It is generally understood that termites lack urate oxidase (uricase), which catalyses the first step of nitrogen recycling from stored uric acid, in their tissues and completely rely on symbiotic microbes in the hindgut for uric acid degradation. Also, termites cannot access the nutritionally useful metabolites produced in the hindgut without trophallaxis (exchange of materials among colony members) [7]. However, most previous research focused only on workers. In contrast, although there is little information regarding uric acid metabolism in reproductive castes, some studies have indicated that termite kings and queens can utilize uric acid by themselves for reproduction [9,16]. These suggest the utilization of uric acid in termites is an ideal model for studying asymmetries in the capability of nitrogen use within insect colonies.

The Japanese subterranean termite *Reticulitermes speratus* (figure 1) is one of the most well-studied termite species, in terms of caste-specific adaptations, due to its unique reproductive system named asexual queen succession (AQS) [17]. As observed in other termite species, *R. speratus* colonies are founded by a monogamous pair of primary reproductives (one king and one queen) derived from alates (winged adults). AQS allows queens to extend their reproductive life because a primary queen (PQ) is genetically immortal until a colony dies due to the production of a large number of secondary queens (SQs) through parthenogenesis [18]. In contrast to queen succession, a primary king (PK) should continue to live even if queens are replaced, because mother-son inbreeding depression can occur when the PK dies and secondary kings (SKs) take over the reproduction duties in the colony [19]. The expression of many caste-specific genes has been reported in this species, in the context of longevity, metabolism and reproduction [20–23]. Therefore, a comparative analysis among castes using *R. speratus* provides an excellent opportunity for investigating asymmetries in the capability of uric acid utilization in termites.

**Figure 1.**
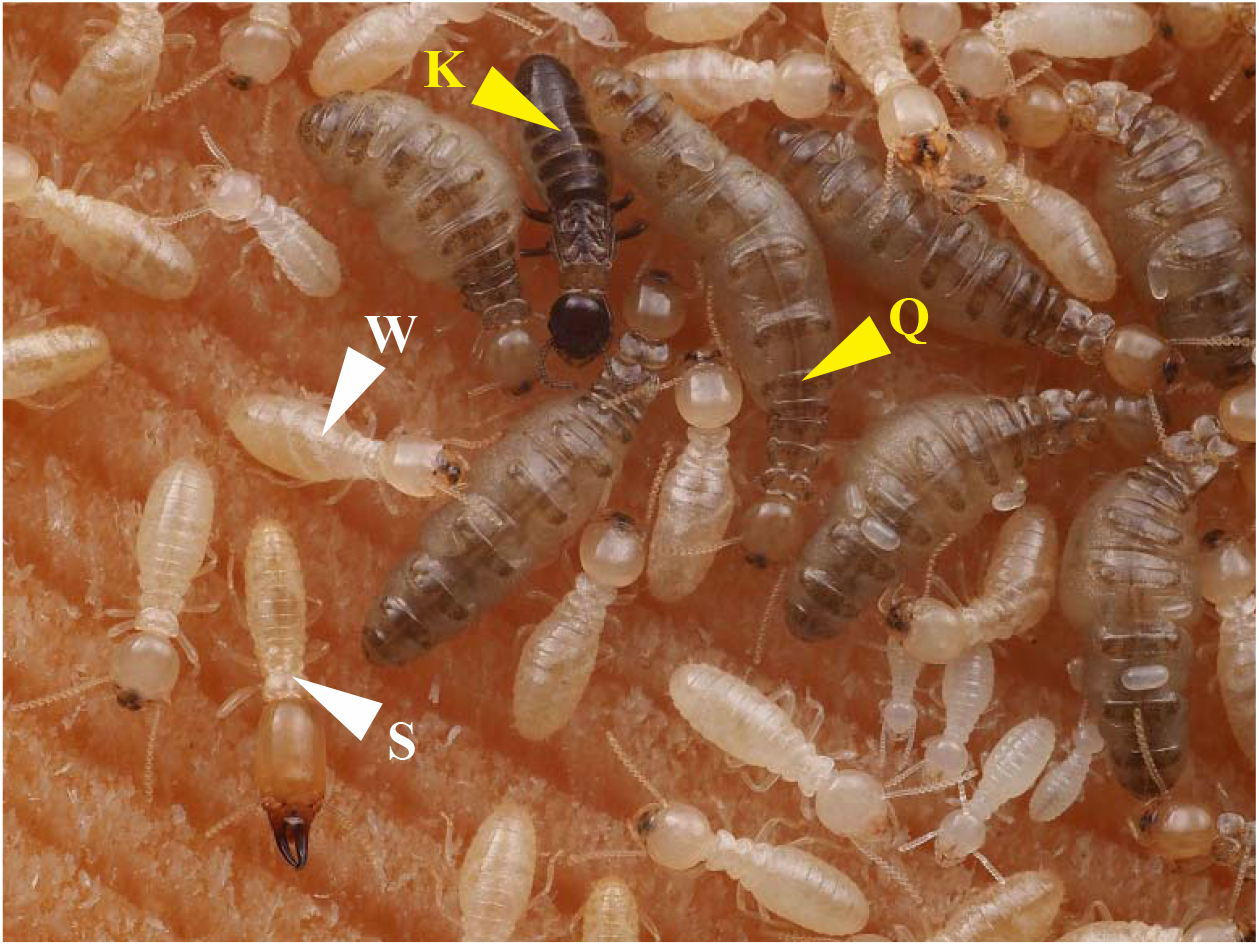
Representative photo of the study termite *Reticulitermes speratus* with each caste. K, Q, W and S refer to king, queen, worker and soldier, respectively. In *R. speratus*, a large number of queens are produced through parthenogenesis. Yellow/white colour indicate reproductive/non-reproductive castes. (Online version in colour.)

In this study, we identified transcripts for enzymes involved in uric acid metabolism (figure 2*a*) from an RNA sequencing (RNA-seq) database of all castes of *R. speratus* (both male and female workers, soldiers, alates, young PKs/PQs and mature PKs/SQs) published in our previous study [20]. Because we found high expression of urate oxidase gene (*RsUAOX*) in mature kings and queens, we additionally examined where *RsUAOX* is overexpressed in the termite body via a caste- and tissue-specific quantitative polymerase chain reaction (qPCR) analysis. To investigate the relationship between uric acid degradation and reproduction, we compared *RsUAOX* expression and uric acid content between workers and worker-derived neotenic kings/queens. In addition, we suppressed uric acid degradation using the RNAi of *RsUAOX* and administration of a urate oxidase inhibitor. Finally, we investigated whether uric acid is provided by workers to reproductive castes within colonies.

**Figure 2.**
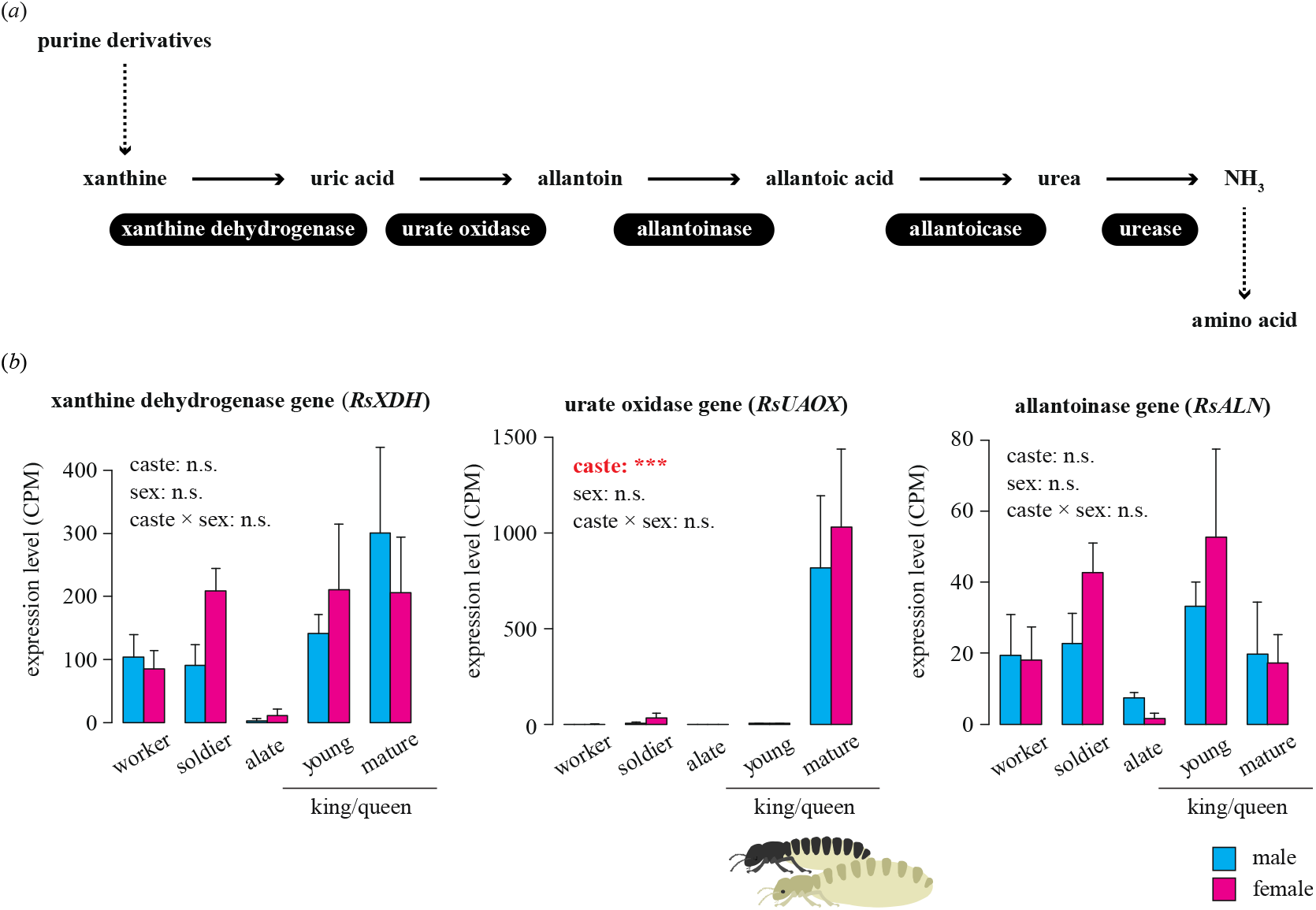
Digital transcriptome analysis of enzyme genes involved in uric acid metabolism using the RNA sequencing (RNA-seq) database of all castes of *Reticulitermes speratus*. (*a*) Generally known pathway of nitrogen recycling in termite workers with relevant enzymes. It has been believed that termites lack urate oxidase in their tissues and completely rely on symbiotic microbes in their hindguts for uric acid degradation. (*b*) Expression patterns of each identified enzyme gene. Comparison of the mean counts per million (CPM) of three transcripts among castes and between sexes. Allantoicase gene and urease gene were not detected in any castes. Young kings/queens refer to young primary kings (PKs) and primary queens (PQs). Mature kings/queens refer to mature PKs and secondary queens (SQs). Cyan and magenta colour indicate male and female, respectively. Error bars represent the standard error of the mean. Results of statistical analysis for each gene expression are shown in the upper left side of each graph [n.s., not significant; *False Discovery Rate (FDR) < 0.05, **FDR < 0.01, ***FDR < 0.001]. (Online version in colour.)

## 2. Materials and methods

### (a) Samples and treatments

Colonies of *R. speratus* were collected along with nest wood from secondary forests in Japan from 2013 to 2021. Due to the rarity of mature PQs in nature [18], we used SQs as mature queens in this study. All colonies were maintained at 25°C in the laboratory and fed mixed sawdust bait [24]. For RNA-seq and qPCR analysis, whole bodies or separated tissues of termites were temporarily preserved at −80°C. The sex of each caste was distinguished by the morphology of caudal sternites following a previous study [25]. No specific permissions were required for these studies.

Firstly, to determine the presence of transcripts for enzymes involved in uric acid metabolism, we performed an RNA-seq analysis in all castes of *R. speratus* [male and female workers, soldiers, alates, young kings/queens (PKs/PQs) and mature kings/queens (PKs/SQs)]. During the swarming season (from April to May), alates were collected from three colonies. Then, five pairs of alates were randomly extracted from each colony and set up in a Petri dish (90 mm in diameter) filled with mixed sawdust bait [24]. After 6 months, they were extracted from each incipient colony and regarded as young kings/queens. Workers, soldiers and mature PKs/SQs were collected from four colonies during the reproductive season (from July to October). Because we observed high expression of urate oxidase gene (*RsUAOX*) in mature kings and queens, we additionally examined where *RsUAOX* is overexpressed in the termite body by caste- and tissue-specific qPCR. For the qPCR analysis, workers, soldiers and mature PKs/SQs were collected from six colonies during the reproductive season.

Next, to investigate the relationship between uric acid degradation and reproduction, comparisons between workers and worker-derived neotenic kings/queens were performed through developmental experiments. Two-hundred workers were set up in a Petri dish (55 mm in diameter) filled with mixed sawdust bait [24]. In such orphaned colonies, some *Reticulitermes* workers are able to differentiate into reproductives called ergatoids, which are recognized on the basis of their slightly heavier cuticle pigmentation, elongated abdomen and wider thorax than workers [26,27]. After 3 months, one male and one female worker, and one neotenic king and one neotenic queen, were randomly extracted. The expression levels of *RsUAOX* (two replicates per colony) and uric acid contents (three replicates for each colony) were compared between workers and neotenic kings/queens. We performed four colony replicates.

In addition, we suppressed uric acid degradation using the RNAi of *RsUAOX*, and by administration of a urate oxidase inhibitor. For RNAi, four colonies (workers and mature queens) were prepared. Small interfering RNA (siRNA) solutions targeting the *RsUAOX* (target gene) were injected into the SQ. Controls were injected with siRNA solution targeting the enhanced green fluorescent protein gene (*EGFP*). An injected queen and 100 workers were set up in a Petri dish (35 mm in diameter) filled with mixed sawdust bait [24]. After 7 days, we performed each injection treatment again. After another 7 days, we counted the eggs laid and conducted qPCR to estimate the efficiency of gene knockdown. The short-term effects of RNAi on target gene expression in individuals sampled 24 h after injection with siRNA were also investigated by qPCR. We performed two replicates for each of the above four colonies. For administration of urate oxidase inhibitor, five colonies (workers and mature queens) were prepared. Urate oxidase inhibitor was injected into the SQ. Controls were injected with just distilled water (DW). An injected queen and 100 workers were set up in a Petri dish (35 mm in diameter) filled with mixed sawdust bait [24]. After 7 days, we counted the number of eggs laid.

Finally, we investigated whether uric acid is provided by non-reproductive castes to reproductive ones. We analysed the queens’ midgut contents, which are composed primarily of trophallactic fluid from workers. Mature kings and queens completely depend on workers for feeding [28]. Uric acid detected in the midgut contents of queens is thought to have originated from workers because uric acid, which is formed in the fat body, does not re-enter the midgut. From each of the five colonies, we randomly selected five mature queens and five workers, and quantified the amount of uric acid in the midgut contents of each individual. The midguts were removed from all individuals by softly holding the termite body and removing the last two abdominal segments using fine-tipped forceps with a soft pulling motion. The midgut contents were then squeezed out of midgut tissues using the forceps. In addition, we examined the effects of royal presence on the uric acid content in worker bodies. The uric acid content was compared among king and queen groups, king-only groups, queen-only groups and royal-absent groups. King and queen groups consisted of a PK, SQ and 500 workers. King-only groups consisted of a PK and 500 workers. Queen-only groups consisted of an SQ and 500 workers. Royal-absent groups consisted of 500 workers. Each group was set up in a plastic box (100 × 100 × 29 mm) filled with mixed sawdust bait [24]. After 3 months, we randomly selected five male and five female workers and quantified the uric acid content in each individual. We performed five colony replicates for each treatment.

### (b) Digital transcriptome analysis of the enzyme genes involved in uric acid metabolism

We analysed a transcriptomic database of all castes of *R. speratus* (male and female workers, soldiers, alates, young kings/queens and mature kings/queens) published in our previous study [20]. After trimming the raw sequencing reads, the remaining reads from all samples were assembled *de novo* using Trinity v2.5.0 [29], which generates transcriptomic assemblies from short read sequences using the de Bruijn graph algorithm. Further details of the assembly and annotation methods are provided in our previous study [20]. Sequence data were deposited in the DNA Data Bank of Japan (DDBJ) under BioProject PRJDB3531, which provides links and access to sample data through the BioSample SAMD00026264–SAMD00026323 and Sequence Read Archive DRR030795–DRR030854. The expression levels of enzyme genes involved in uric acid metabolism were estimated using RSEM v1.2.8 [30] for the filtered reads from each sample. Raw read counts generated by RSEM were normalized using the trimmed mean of M-value (TMM) normalization method [31]. The read counts were then used for differential expression analyses, among castes and between sexes, using the “edgeR” package (v3.4.2) [32] in R software (v4.0.5) [33]. We confirmed the presence of the urate oxidase gene *RsUAOX* by performing a phylogenetic analysis (see electronic supplementary material, Text S1).

### (c) qPCR analysis of the urate oxidase gene

The urate oxidase homolog of *R. speratus* (*RsUAOX*; accession number pending) was obtained from the RNA sequencing database published in our previous study [20]. through a basic local alignment search tool (BLAST) search with the amino acid sequences of translated urate oxidase genes in some insect species. We designed primer pairs for *RsUAOX* using Primer3Plus (https://primer3plus.com/cgi-bin/dev/primer3plus.cgi). Five reference primers for glucose-6-phosphate 1-dehydrogenase (*RsG6PD*; accession number: FX983730), glyceraldehyde-3-phosphate dehydrogenase (*RsGAPDH*; accession number: FX983172), beta-actin (*RsACT*; accession number: FX983744), NADH dehydrogenase subunit 5 (ND5) and elongation factor-1 alpha (EF1α) were referred to from a previous study [34,35]. The primer sequences are shown in the electronic supplemental material (table S2). To select the best control gene, the suitability of the genes of this species was evaluated using NormFinder software [36]. Using an RNeasy mini kit (Qiagen, Hilden, Germany), total RNA was extracted individually from the whole bodies or separated tissues (fat bodies, testes/ovaries and other tissues) of termites prepared by the above method. For the tissues of workers and soldiers, we pooled three individuals to extract a sufficient amount of RNA for analysis. The cDNA was synthesized immediately from the RNA using a PrimeScript™ RT reagent kit (Takara Bio, Shiga, Japan) and preserved at −20°C for the qPCR analysis. The gene expression levels were determined by qPCR performed using an Applied Biosystems^®^ StepOne™ system with Power SYBR™ Green PCR master mix (Thermo Fisher Scientific, Waltham, MA, USA). All procedures were performed in accordance with the manufacturer's protocol. Relative expression levels were calculated using a typical 2^(−ΔΔ*C*_T_) method [37].

### (d) Quantification of uric acid content

Uric acid concentrations were measured using a Uric Acid Assay Kit (Abcam, Cambridge, UK) according to the manufacturer’s protocols. Briefly, each sample was homogenized in a uric acid assay buffer and centrifuged at 15,000 rpm for 2 min at 4°C. Supernatants were collected as sample extracts, and a reaction mix was added for 30 min at 37°C in the dark. The uric acid content was determined fluorometrically (Ex/Em = 535/587 nm) and compared with serially diluted uric acid standards (0, 0.8, 1.6, 2.4, 3.2, and 4.0 nmol/well). To eliminate differences in body size, the uric acid content was standardised by dividing by the fresh body weight.

### (e) Suppression of uric acid degradation

The RNAi of *RsUAOX* was administered via injection. The siRNA were designed using siDirect software (v2.0; http://sidirect2.rnai.jp/). Two siRNAs were designed and mixed (100 μM each) to target specific sequences of the *RsUAOX* common region (*RsUAOX*_1, sense: 5’-CGACAAUAAGGACAUCAUUGC-3’, antisense: 5’-AAUGAUGUCCUUAUUGUCGCC-3’; *RsUAOX*_2, sense: 5’-CAUAACCACGCCUUCAUCUUC-3’, antisense: 5’-AGAUGAAGGCGUGGUUAUGAU-3’). As in previous studies [38,39], the enhanced green fluorescent protein (*EGFP*) gene was selected as a control gene (*EGFP*, sense: 5’-GCAUCAAGGUGAACUUCAAGA-3’, antisense: 5’-UUGAAGUUCACCUUGAUGCCG-3’). One microlitre of 100 μM siRNA solution was injected into the ventral side of the mature queen’s thorax using a FemtoJet^®^ 4i microinjector (Eppendorf, Hamburg, Germany) fitted with a custom-pulled glass capillary (Narishige, Tokyo, Japan), with double-sided tape used to facilitate termite immobilization. In addition, urate oxidase inhibitor was injected. Potassium oxonate is a potent in vivo inhibitor of urate oxidase [40]. One microlitre of 10 mM potassium oxonate was injected into the ventral side of the mature queen’s thorax using a FemtoJet^®^ 4i microinjector (Eppendorf) fitted with a custom-pulled glass capillary (Narishige), with double-sided tape used to facilitated termite immobilization. Controls were injected with an equivalent volume of DW. The methods used were modified versions of those used in a rat study (*Rattus norvegicus*) [40].

### (f) Data analysis

All statistical analyses were performed using R software (v4.0.5) [33]. We analysed the relative gene expression levels and uric acid contents using linear models (LMs) and linear mixed models (LMMs). In the LM, treatment was a fixed factor. In the LMM, treatment was a fixed factor and colony was a random factor. Likelihood ratio tests were conducted to determine the significance of each fixed effect, and Tukey's HSD tests were conducted to test for differences between the levels of a significant fixed effect. We analysed data on the number of eggs laid using generalized linear models (GLMs) with a Poisson distribution. In the GLMs, treatment was a fixed factor. Likelihood ratio tests were conducted to test the significance of each fixed effect. All data in the figures are presented as the mean + standard error (SE). Asterisks indicate significant differences at \**p* < 0.05, \*\**p* < 0.01, \*\*\**p* < 0.001. Different letters over the bars indicate significant differences at *p* < 0.05.

## 3. Results

### (a) Digital transcriptome analysis of the enzyme genes involved in uric acid metabolism

Using the RNA-seq database of all castes of *R. speratus*, we identified three enzyme genes involved in uric acid metabolism [xanthine dehydrogenase gene (*RsXDH*), urate oxidase gene (*RsUAOX*) and allantoinase gene (*RsALN*); figure 2*b*]. Notably, mature kings and queens had higher expression levels than the other castes of *RsUAOX*, which catalyses the oxidation of uric acid [caste: false discovery rate (FDR) < 0.001, sex: FDR = 1.000, caste × sex: FDR = 1.000; figure 2*b*]. No significant differences were observed, among castes or between sexes, in the expression levels of *RsXDH* (caste: FDR = 1.000, sex: FDR = 1.000, caste × sex: FDR = 1.000; figure 2*b*) and *RsALN* (caste: FDR = 0.557, sex: FDR = 1.000, caste × sex: FDR = 1.000; figure 2*b*). In this analysis, the allantoicase and urease genes were not detected in any castes.

### (b) Caste- and tissue-specific qPCR analysis of *RsUAOX*

The fat bodies of mature kings and queens had high expression levels of *RsUAOX*. The combination of caste (worker, soldier or reproductive), tissue (fat body, testis/ovary or other tissues) and sex (male or female) had a significant effect on the expression level of *RsUAOX* (the interaction term among caste, tissue and sex was included; likelihood ratio test, *df* = 13, χ^2^ = 114.56, *p* < 0.001, figure 3). The relative *RsUAOX* expression levels in the fat bodies of reproductives were significantly higher than those of workers (Tukey’s HSD test, male: *p* < 0.001; female: *p* < 0.001, figure 3) and soldiers (Tukey’s HSD test, male: *p* < 0.001; female: *p* < 0.001, figure 3). This trend was seen for both males and females. No significant differences were observed between males and females in the relative expression levels of *RsUAOX* in fat bodies (Tukey’s HSD test, worker: *p* = 1.000, soldier: *p* = 1.000, reproductive: *p* = 1.000, figure 3). Expression levels relative to fat bodies of male workers are shown in the figure.

**Figure 3.**
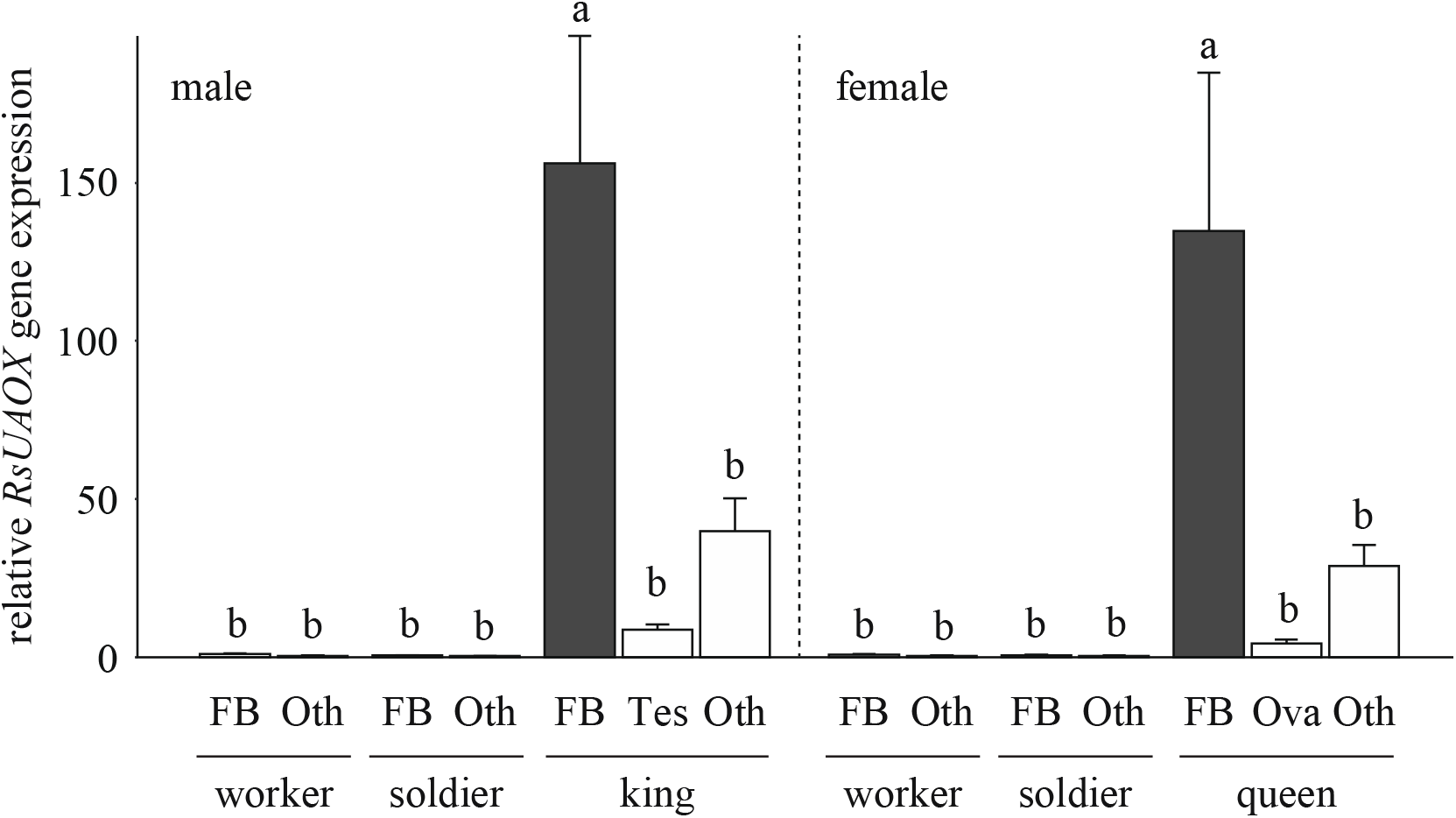
Caste- and tissue-specific quantitative PCR (qPCR) analysis of urate oxidase gene (*RsUAOX*). Comparison of the relative expression levels of *RsUAOX* among castes and tissues. Expression levels relative to fat bodies of male workers are shown. Kings/queens refer to mature primary kings (PKs) and mature secondary queens (SQs). Error bars represent the standard error of the mean. Different letters indicate significant differences (Tukey’s HSD tests, *p* < 0.05). FB, fat body; Tes/Ova; testis/ovary; Oth; the others.

### (c) Developmental experiments

Intriguingly, worker-derived neotenic kings and queens also had high expression levels of *RsUAOX*. The combination of caste (worker or neotenic) and sex (male or female) had a significant effect on the expression level of *RsUAOX* (the interaction term between caste and sex was included; likelihood ratio test, *df* = 3, χ^2^ = 25.12, *p* < 0.001, figure 4*a*). The relative *RsUAOX* gene expression levels of neotenic kings/queens were significantly higher than those of workers (Tukey’s HSD test, male: *p* = 0.00966; female: *p* < 0.001, figure 4*a*). Expression levels relative to male workers are shown in the figure. In addition, neotenic kings/queens had low uric acid contents. The combination of caste (worker or neotenic) and sex (male or female) had a significant effect on uric acid content (the interaction term between caste and sex was included; likelihood ratio test, *df* = 3, χ^2^ = 41.52, *p* < 0.001, figure 4*b*). The uric acid contents of neotenic kings/queens were significantly lower than those of workers (Tukey’s HSD test, male: *p* = 0.0146; female: *p* < 0.001, figure 4*b*). This trend was seen for both males and females. No significant differences were observed between males and females in the relative expression levels of *RsUAOX* (Tukey’s HSD test, neotenic: *p* = 0.867; worker: *p* = 1.000, figure 4*a*) or the uric acid contents (Tukey’s HSD test, neotenic: *p* = 0.337; worker: *p* = 0.736, figure 4*b*).

**Figure 4.**
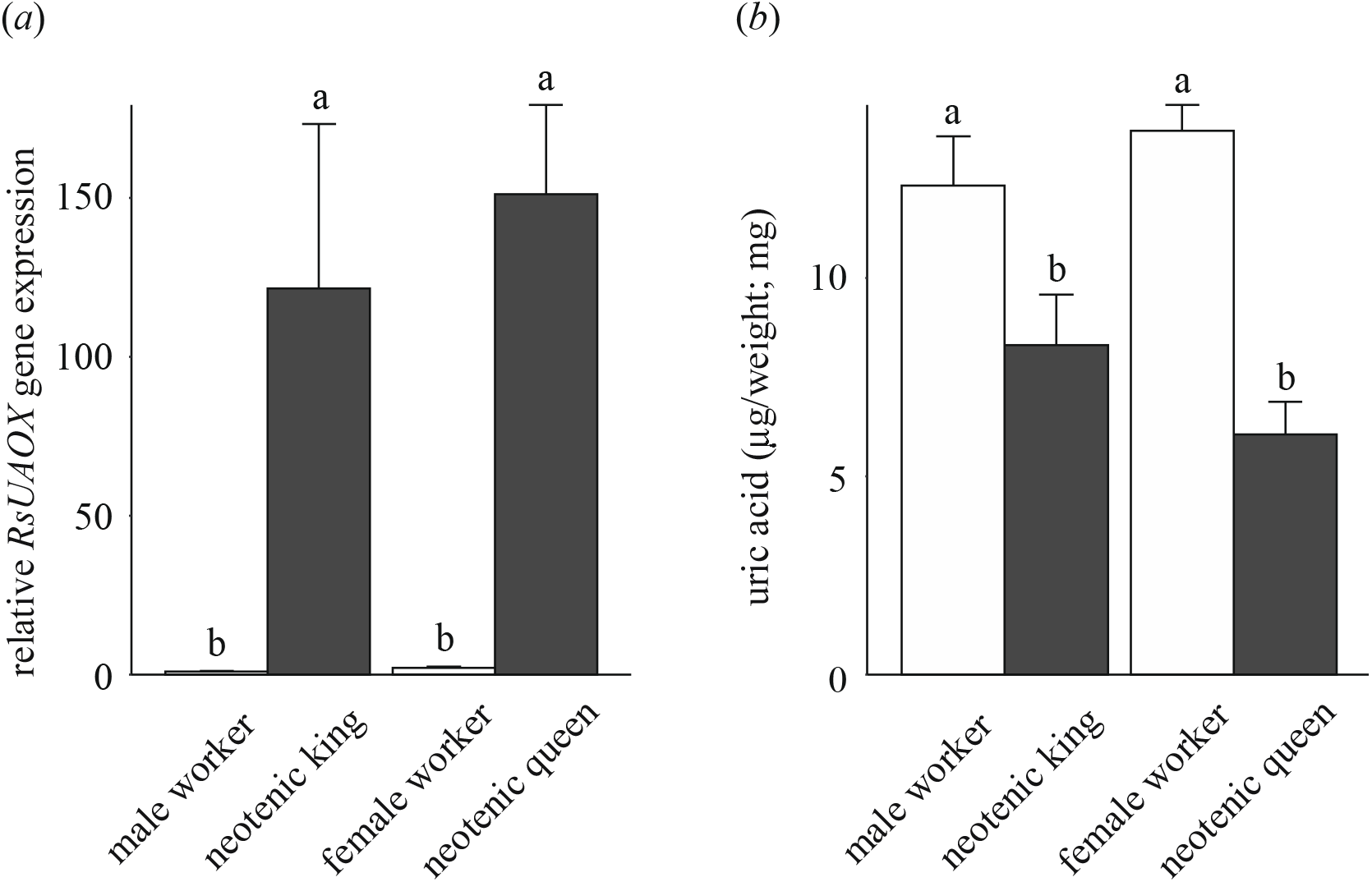
Comparison between workers and worker-derived neotenic kings/queens performed during developmental experiments. In royal-absent conditions, some *Reticulitermes speratus* workers are able to differentiate into reproductives. (*a*) Comparison of the relative expression levels of urate oxidase gene (*RsUAOX*) by quantitative PCR (qPCR). Expression levels relative to male workers are shown. (*b*) Comparison of the uric acid contents. Error bars represent the standard error of the mean. Different letters indicate significant differences (Tukey’s HSD tests, *p* < 0.05).

### (d) Suppression of uric acid degradation

The effects of injection of the RNAi of *RsUAOX* (target gene) and *EGFP* (control gene) were investigated. *RsUAOX* knockdown significantly decreased the number of eggs laid per queen (likelihood ratio test, *df* = 1, χ^2^ = 10.90, *p* < 0.001, figure 5*a*), while there were no significant differences between the *RsUAOX* and *EGFP* treatments in the expression levels of the target gene (likelihood ratio test, *df* = 1, χ^2^ = 0.51, *p* = 0.475). The short-term effects of RNAi on target gene expression were also investigated by qPCR in individuals sampled 24 h after injection with siRNA. The relative *RsUAOX* gene expression levels of *RsUAOX* were significantly lower than those of *EGFP* (likelihood ratio test, *df* = 1, χ^2^ = 5.16, *p* = 0.0317, figure 5*b*). Thus, RNAi is effective, although its functional period in termites is short. We also confirmed that there were no significant differences between the two treatments in the initial body weights of queens (likelihood ratio test, long-term: *df* = 1, χ^2^ = 0.38, *p* = 0.472; short-term: *df* = 1, χ^2^ = 0.017, *p* = 0.892). Expression levels relative to *EGFP* are shown in the figure. In addition, the effects of injection of potassium oxonate (urate oxidase inhibitor) and DW (control) were investigated. Administration of the urate oxidase inhibitor significantly decreased the number of eggs laid per queen (likelihood ratio test, *df* = 1, χ^2^ = 27.93, *p* < 0.001, figure 5*c*). We also confirmed that there were no significant differences between the two treatments in the initial body weights of queens (likelihood ratio test, *df* = 1, χ^2^ = 0.13, *p* = 0.718).

**Figure 5.**
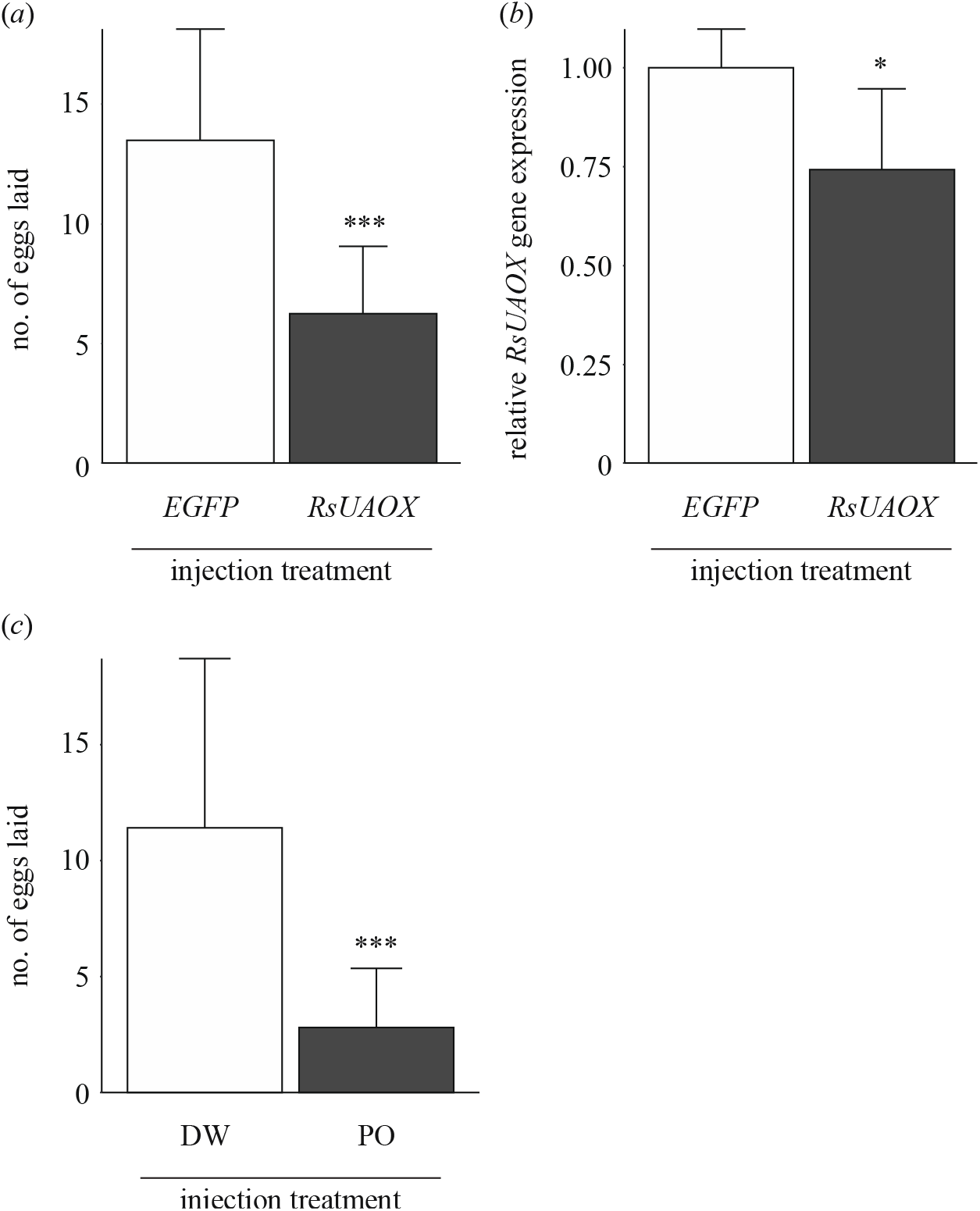
Effects of suppression of uric acid degradation on reproductive activity in termites. RNA interference (RNAi) of urate oxidase gene (*RsUAOX*) and administration of urate oxidase inhibitor were performed. (*a*) Effects of gene knockdown of *EGFP* (control gene) and *RsUAOX* (target gene) on egg production in termite queens. The number of eggs was compared between treatments. (*b*) Short-term (24 h after injection) effects of gene knockdown of *EGFP* and *RsUAOX* on gene expression. The relative expression levels of *RsUAOX* by quantitative PCR (qPCR) were compared between treatments. Expression levels relative to *EGFP* are shown. (*c*) Effects of administration of distilled water (DW; control) and potassium oxonate (PO; urate oxidase inhibitor) on egg production in termite queens. The number of eggs was compared between treatments. Error bars represent the standard error of the mean. Asterisks indicate significant differences (likelihood ratio test, \**p* < 0.05, \*\**p* < 0.01, \*\*\**p* < 0.001).

### (e) Exchange of uric acid within colonies

Uric acid was detected in the contents of the queens’ midgut (figure 6*a*), where trophallactic fluid from workers is harboured. The uric acid content in the midguts of queens was significantly higher than in the midguts of workers (likelihood ratio test, *df* = 1, χ^2^ = 23.38, *p* < 0.001, figure 6*b*). Intriguingly, the absence of kings or queens enhanced uric acid accumulation in worker bodies. The combination of royal presence (king and queen group, king-only group, queen-only group or royal-absent group) and sex (male or female) had a significant effect on uric acid content (the interaction term between royal presence and sex was included; likelihood ratio test, *df* = 7, χ^2^ = 101.43, *p* < 0.001, figure 6*c*). The uric acid contents of workers in the king and queen groups were significantly lower than in the workers in king-only groups (Tukey’s HSD test, male: *p* < 0.001; female: *p* < 0.001, figure 6*c*), queen-only groups (Tukey’s HSD test, male: *p* < 0.001; female: *p* < 0.001, figure 6*c*) and royal-absent groups (Tukey’s HSD test, male: *p* < 0.001; female: *p* < 0.001, figure 6*c*). This trend was seen for both males and females. No significant differences in uric acid content were observed between males and females (Tukey’s HSD test, king and queen groups: *p* = 1.000; king-only groups: *p* = 0.960; queen-only groups: *p* = 0.998; royal-absent groups: *p* = 0.999, figure 6*c*).

**Figure 6.**
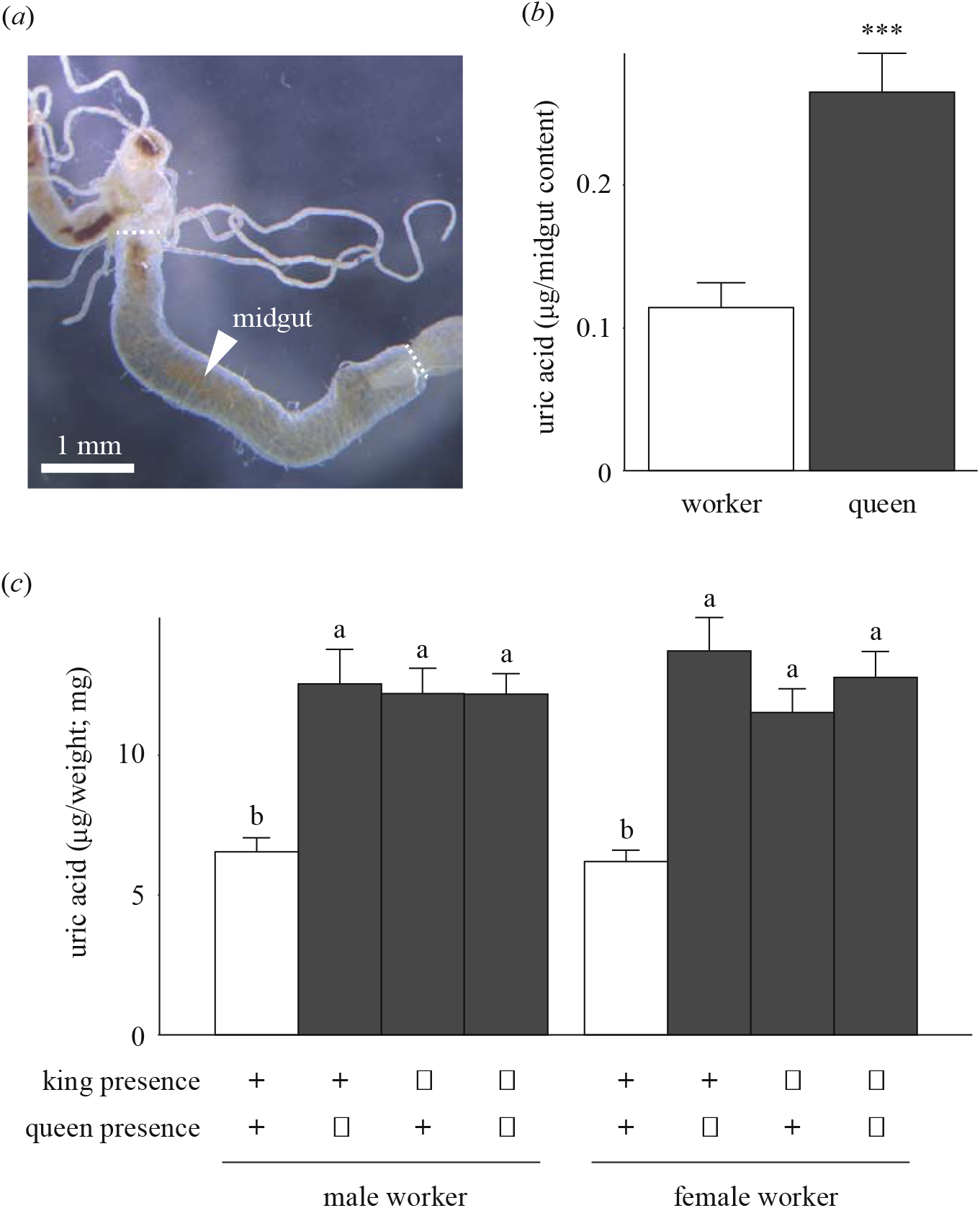
Influences of royal presence on the exchange of uric acid. (*a*) Representative photo of the midgut of the termite queen (indicated by the white arrowhead), where trophallactic fluid from workers is harboured. (*b*) Comparison of the amount of uric acid in midgut contents between workers (non-reproductive caste) and queens (reproductive caste). Asterisks indicate significant differences (likelihood ratio test, \**p* < 0.05, \*\**p* < 0.01, \*\*\**p* < 0.001). (*c*) Comparison of the uric acid contents in worker bodies among king and queen groups, king-only groups, queen-only groups and royal-absent groups under laboratory conditions. Different letters indicate significant differences (Tukey’s HSD tests, *p* < 0.05). Error bars represent the standard error of the mean.

## 4. Discussion

We have shown that the urate oxidase gene *RsUAOX*, which catalyses the first step of nitrogen recycling from stored uric acid, is highly expressed in mature kings and queens compared with the other castes in the subterranean termite *R. speratus*. This study is the first to clearly demonstrate the existence of urate oxidase in termites. Previously, it was believed that termites lack this enzyme in their tissues, instead completely relying on symbiotic microbes in their hindguts for uric acid degradation [8,12–15]. We also showed that uric acid degradation by urate oxidase is the key to reproduction in termites. Differentiation from totipotent workers into neotenic kings/queens was accompanied by increased expression of *RsUAOX* and decreased amounts of uric acid. Suppression of uric acid degradation by the RNAi of *RsUAOX*, and administration of a urate oxidase inhibitor, decreased the number of eggs laid per queen. Uric acid was revealed to be provided from workers to reproductive castes via trophallaxis. These results strongly support our hypothesis of an asymmetry in the capacity for uric acid utilization; termite kings and queens can degrade uric acid by themselves to aid reproduction, while the other non-reproductive castes cannot. Furthermore, in the dampwood termite *Zootermopsis nevadensis*, a previous study reported that feeding on a diet supplemented with uric acid promotes offspring production by kings and queens [9]. According to Elliott and Stay [16], the fat bodies of physogastric (with an enlarged abdomen) queens of *R. flavipes* contain fewer urocyto cells than non-physogastric queens. These authors related this observation to the utilization of the nitrogen of these urates for the intense production of eggs in physogastric queens.

The utilization of stored uric acid in insects has often been discussed in terms of the metabolic pathway shared with endosymbionts. For example, in the brown planthopper *Nilaparvata lugens*, uric acid utilization has been attributed to eukaryotic yeast-like endosymbionts [41]. In the shield bug *Parastrachia japonensis*, the presence of *Erwinia*-like bacteria localized in midguts has been demonstrated to be responsible for enzymatic uricolysis [42]. Cockroaches harbour intracellular bacteria within their fat bodies, named *Blattabacterium*, which participate in the nitrogen recycling process [43,44]. Although termites are close relatives of cockroaches, *Blattabacterium* has been lost in all species except the basal termite species *Mastotermes darwiniensis* [11]. Instead, in termites, it is believed that anaerobic bacteria in the hindguts enzymatically liberate uric acid nitrogen into a usable form (i.e. ammonia) [8,12–15]. However, termites cannot access the nutritionally useful metabolites produced in the hindguts without trophallaxis [7]. In this study, we demonstrated that only kings and queens can degrade uric acid with their own enzymes in fat bodies, resulting in increased reproductive output, while genes of the downstream enzymes that independently synthesize ammonia (i.e. allantoicase and urease) were not detected. We therefore believe that the downstream pathway of uric acid utilization in termite kings and queens is different from the known pathway of nitrogen recycling in workers. Future studies should directly investigate the metabolic process from stored uric acid to reproductive output in termites using ^15^N-labeled uric acid. Also, in the study termite *R. speratus*, whereas workers and soldiers have an abundance of protists in their hindguts, kings and queens have no protists [45]. This asymmetry in the abundance of symbiotic microbes may be related to the royal-specific degradation of urate oxidase.

Why do termite kings and queens degrade uric acid by themselves in their fat bodies, instead of directly receiving useful downstream metabolites from workers? It is possible that uric acid is the best form to exchange due to its chemical properties (e.g. the low solubility of uric acid and its salts in water can reduce water loss from individuals during trophallaxis). It is also possible that uric acid degradation is not only undertaken for acquiring a nitrogen source, but has another important function for termite kings and queens. Investigating the biological functions of uric acid and its derivatives in termites is therefore of interest. For example, uric acid has been demonstrated to be an antioxidant against reactive oxygen species (ROS), which can cause damage to biomolecules such as DNA, proteins and lipids in termites [46]. Also, uric acid is enzymatically oxidized by urate oxidase to produce allantoin, hydrogen peroxide (H_2_O_2_) and carbon dioxide (CO_2_). Allantoin is known to promote cell proliferation, wound healing and stress tolerance in animals and plants [47,48]. According to Shestopalov et al. [47], in humans the allantoin level is increased in the placenta in weeks 7–8 of pregnancy, which represents the period of the first wave of cytotrophoblast invasion into the endometrium and is accompanied by active cell proliferative processes. The authors proposed that allantoin plays an important role in these events. Long-term high reproductive outputs of termite kings and queens may be due to the functions of allantoin [49].

Royal presence is crucial to the exchange of uric acid within colonies. In this study, the absence of kings/queens, which are recipients of uric acid, enhanced uric acid accumulation in worker bodies. Interestingly, no difference was observed between the effects of losing both versus one reproductive. There may be a maximum amount of uric acid that can be accumulated in a worker’s body. Considering the importance of uric acid degradation in termite reproduction, it is reasonable to suggest that totipotent workers can acquire resources for reproduction prior to the differentiation into neotenic kings/queens triggered by royal-absence. Our results were consistent with those of previous studies reporting that termite workers gradually accumulate uric acid inside fat bodies after maintenance under laboratory conditions (i.e. separated from royal chambers) for several months [12,50,51]. The absence of a particular caste, especially kings and queens, within the colony causes various physiological and behavioural changes in termites [52–55]. This study provides further empirical evidence that taking account of the social context is important when conducting ecological studies using termites.

In conclusion, the enzymatic degradation of uric acid specific to mature kings and queens is the key to reproduction in termites. Our results suggest that the capacity for nitrogen use, which is essential for the protein synthesis required for reproduction, at least partially maintains colony cohesion expressed as the reproductive monopoly held by the kings and queens. This study sheds light on the nature of the reproductive division of labour in social insects from the perspective of resource use. Furthermore, considering applied biology, termites are one of the most economically important pests in the world, and one of the most difficult to control. Our findings may lead to future technical innovations in termite control, such as the discovery of inhibitors that can supress reproductive activity.

## Supporting information

electronic supplementary material

## Data accessibility

The dataset is available as electric supplementary material (dataset S1).

## Authors’ contributions

TK: conceptualization, data curation, formal analysis, funding acquisition, investigation, methodology, resources, validation, visualization, writing-original draft; ET: conceptualization, data curation, funding acquisition, investigation, methodology, resources, visualization, writing-review and editing; MT: conceptualization, funding acquisition, methodology, resources, visualization, writing-review and editing; KM: conceptualization, funding acquisition, project administration, resources, supervision, writing-review and editing. All authors gave final approval for publication and agreed to be held accountable for the work performed therein.

## Acknowledgements

We thank Tomoki Ishibashi, Shuya Nagai, Takehiro Morimoto, Chihiro Tamaki, Hiroki Noda and Hikaru Mitomo for their assistance in collecting termites; Yoshihito Iuchi, Yuki Mitaka, Michihiko Takahashi, and Matthew Kamiyama for helpful discussion; Soshi Araki for photographs; and Shigeto Dobata and Kazuya Kobayashi for their assistance with data analysis. We also thank Kunpeng Liu, Yao Wu, Shun Mizote and other members of the Laboratory of Insect Ecology, Kyoto University for inspiring scientific discussions.

