## Supplementary material for "King- and queen-specific degradation of uric acid is the key to reproduction in termites": electronic supplementary material

**This file includes:**

Text S1: Supplementary Materials and Methods

Figure S1

Table S1 and S2

**Other Supplementary Materials for this manuscript includes the following:**

Dataset S1

**Text S1: Supplementary Materials and Methods**

**Phylogenetic analysis of urate oxidase (uricase) homolog sequences in insects**

To confirm the identities of the urate oxidase gene *RsUAOX*, we performed multiple amino acid sequence alignments with MUSCLE, trimmed the alignment sequences using trimAl version 1.2rev59 [1], and conducted phylogenetic analyses using the molecular evolutionary genetics analysis software MEGA X [2] (figure S1). Gene evolutionary history was inferred using the maximum likelihood method based on the Le_Gascuel_2008 model [3], which is the best model based on the Bayesian information criterion. Urate oxidase homologs of termite species (*R. speratus* and *Cryptotermes secundus*) and the other insects (Blattodea, Hemiptera, Hymenoptera, Coleoptera, Lepidoptera and Diptera) were analysed. Furthermore, analyses of protein families and domains were also performed using InterProScan (https://www.ebi.ac.uk/interpro/) (table S1). The gene *RsUAOX* sequence contains the conserved site and domains of urate oxidase genes.

**Figure S1. Maximum likelihood molecular phylogenetic tree of urate oxidase homolog sequences**

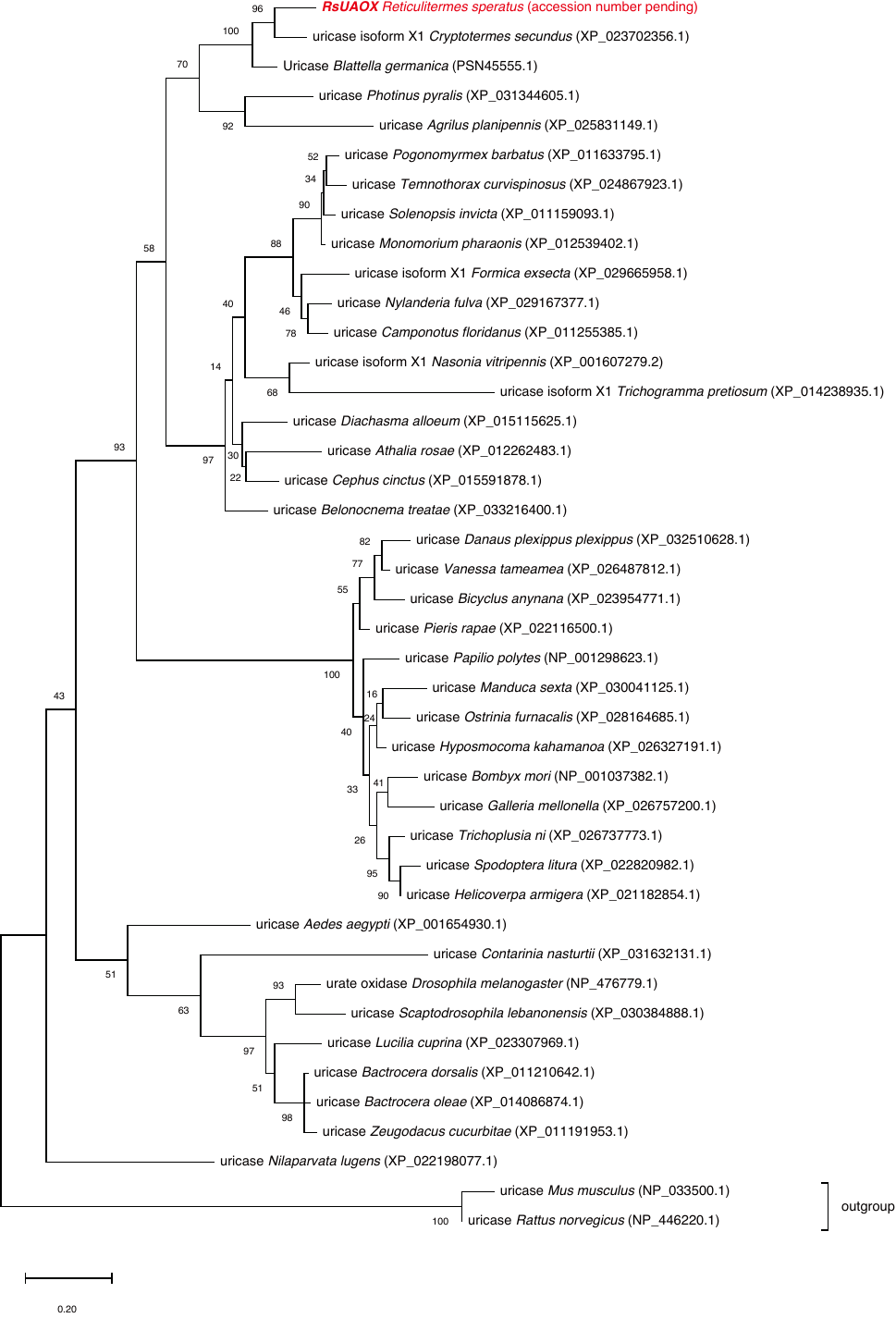

**Figure S1. Maximum likelihood molecular phylogenetic tree of urate oxidase homolog sequences**

The evolutionary history of urate oxidase homologs was inferred using the maximum likelihood method based on the Le_Gascuel_2008 model [3]. The tree with the highest log likelihood (-6564.33) is shown. The percentage of trees in which the associated taxa clustered together is shown next to the branches. Initial tree(s) for the heuristic search were obtained automatically by applying Neighbor-Join and BioNJ algorithms to a matrix of pairwise distances estimated using the JTT model, and then selecting the topology with superior log likelihood value. A discrete Gamma distribution was used to model evolutionary rate differences among sites [five categories (+*G*, parameter = 0.5936)]. The tree is drawn to scale, with branch lengths measured in the number of substitutions per site. This analysis involved 42 amino acid sequences. There was a total of 258 positions in the final dataset. Evolutionary analyses were conducted in MEGA X [2].

**Table S1. Predicted *RsUAOX* families and domains**

| Gene name | AC (sequence) | Type | Name | GO IDs | Library |
| --- | --- | --- | --- | --- | --- |
| *RsUAOX* | IPR002042  (18–326) | Domain | Uricase | - | TIGR03383 (TIGRFAMs)  PF01014 (PFAM)  PIRSF000241 (PIRSF)  PR00093 (PRINTS)  PTHR42874 (PANTHER) |
|  | IPR019842  (176–203) | Domain | Uricase, conserved site | GO:0006144  GO:0004846 | PS00366 (PROSITE_PATTERNS) |
|  | no IPR  (32–161,  167–319) | Homologous superfamily | Tetrahydrobiopterin biosynthesis enzymes-like | - | SSF55620 (SUPERFAMILY) |
|  | no IPR  (18–324) | Homologous superfamily | Urate Oxidase | - | G3DSA:3.10.270.10 (CATH-Gene3D) |
|  | no IPR  (1–25) | - | Disorder_prediction | - | mobidb-lite (MOBIDB_LITE) |

**Table S2. Primer sequences**

| Target gene | Sequence (5’-3’) | Amplicon (bp) |
| --- | --- | --- |
| *RsUAOX* | Forward; CGTGGTTGAGGCGTCTGTT  Reverse; GAGAGAAGATGAAGGCGTGGTT | 92 |
| *RsND5* | Forward; GCTGGGGGGGTTATTCATTCCAT  Reverse; GGCATACCACAAAGGGCAAAA | 125 |
| *RsEF1α* | Forward; GGTGATGCGGCTATTGTTAACC  Reverse; GTGGTGGGAATTCTGAGAAAGATT | 73 |
| *RsG6PD* | Forward; GCACTTTGTTCGTTCTGATGAGTT  Reverse; TCACTACACACTTCATCTGCCTTCT | 144 |
| *RsGAPDH* | Forward; CCATAGAAAAGGCTTCTGCACATT  Reverse; AACAACAAACATTGGGGCATC | 89 |
| *RsACT* | Forward; AAATCGTGCGTGACATCAAA  Reverse; GGAACAGAGCCTCAGGACAG | 168 |
